# EnrichSci: Transcript-guided Targeted Cell Enrichment for Scalable Single-Cell RNA Sequencing

**DOI:** 10.1101/2025.05.02.651937

**Authors:** Andrew Liao, Zehao Zhang, Andras Sziraki, Abdulraouf Abdulraouf, Zihan Xu, Ziyu Lu, Wei Zhou, Junyue Cao

**Author notes:** Correspondence (W.Z.), (J.C.). Senior author (Wei Zhou, Junyue Cao).

## Abstract

Large-scale single-cell atlas efforts have revealed many aging-or disease-associated cell types, yet these populations are often underrepresented in heterogeneous tissues, limiting detailed molecular and dynamic analyses. To address this, we developed EnrichSci—a highly scalable, microfluidics-free platform that combines Hybridization Chain Reaction RNA FISH with combinatorial indexing to profile single -nucleus transcriptomes of targeted cell types with full gene-body coverage. When applied to profile oligodendrocytes in the aging mouse brain, EnrichSci uncovered aging-associated molecular dynamics across distinct oligodendrocyte subtypes, revealing both shared and subtype-specific gene expression changes. Additionally, we identified aging-associated exon-level signatures that are missed by conventional gene-level analyses, highlighting post-transcriptional regulation as a critical dimension of cell-state dynamics in aging. By coupling transcript-guided enrichment with a scalable sequencing workflow, EnrichSci provides a versatile approach to decode dynamic regulatory landscapes in diverse cell types from complex tissues.

## Main

Mammalian organs are maintained through the homeostasis of hundreds to thousands of distinct cell states, each varying in proportion from common populations like hepatocytes (∼70% of liver cells) to rare types such as pinealocytes (<0.01% of brain cells)^1,2^. While scalable single-cell genomics studies have cataloged many of these populations, the high disparity in their frequencies often leaves rare cell types underrepresented. This under-sampling hinders our ability to characterize the molecular heterogeneity of these cell types and uncover their detailed changes in various conditions, thereby complicating the development of targeted therapeutic interventions.

To tackle these challenges, several groups recently developed methods that couple transcript-based labeling of specific cell types with cell sorting and downstream transcriptome analysis^3^. These techniques bypass the cost and technical constraints of antibody-based enrichment to profile specific cell types efficiently, but each still faces limitations. Probe-seq^4^ first integrated a sensitive RNA FISH method with a downstream bulk RNA-seq workflow to enable the antibody-free isolation and profiling of target cell types, inspiring the development of further techniques to achieve similar targeted profiling at single-cell resolution. FIND-seq^5^ enabled transcript-based cell type-targeted scRNA-seq by coupling PCR-based detection of target transcripts with microfluidic cytometry, but it has limited throughput (up to ∼10^3^ cells) and requires specialized equipment. PERFF-seq^3^, developed most recently, effectively combined RNA FISH-based FACS with a higher throughput scRNA-seq approach (up to ∼10^5^ cells). However, this method utilizes the commercial 10x Flex scRNA-seq platform, which captures a limited view of the transcriptome due to its reliance on a pre-defined probe set, hindering the analysis of samples from different species as well as the dynamics of genome-wide RNA elements (both exonic and intronic expression as well as many non-protein coding genes) that are crucial for cellular state regulation. These trade-offs limit both broad application and comprehensive resolution of rare cell subtypes across large sample sets.

To address the above challenges, we developed EnrichSci—a highly scalable single-cell combinatorial indexing approach^6^ designed for genome-wide analysis of exonic and intronic expression in enriched cell populations. EnrichSci combines our optimized single-cell combinatorial indexing platform EasySci^1^, which can process tens of millions of cells per study at under $0.001 per cell and supports both cellular and nuclear fixation, with a Hybridization Chain Reaction (HCR) RNA FISH workflow^7,8^. This integration enables targeted cell enrichment and sequencing of rare cell types from complex tissues (*e*.*g*., brain), allowing detailed analyses of subtype-specific transcriptional signatures and dynamics across diverse biological conditions.

The EnrichSci workflow begins by selecting a gene module uniquely expressed in the target cell type (**Fig. 1a**), which improves labeling specificity over traditional single-transcript approaches. Next, nuclei are extracted from complex tissues or cell lines, fixed in formaldehyde, and fluorescently labeled using Hybridization Chain Reaction (HCR)^7,8^ RNA FISH before enrichment via FACS (**Fig. 1a**). The enriched nuclei are then processed through a slightly modified EasySci pipeline for single-cell transcriptome profiling (**Fig. 1a**). Notably, by employing both oligo-dT and random hexamer primers during reverse transcription, this approach delivers comprehensive coverage of the entire gene body, including both exons and introns. Our plate-based combinatorial indexing approach also avoids the reliance on custom or commercial microfluidic systems required by other methods^3,5^ and is fully compatible with formaldehyde-fixed nuclei from complex tissues. Moreover, the enrichment pipeline has been extensively optimized (*e*.*g*., lysis buffer, probe concentration, washing conditions) to enhance signal specificity of the RNA FISH workflow and improve cell recovery rates (**Extended Data Fig. 1a-c**).

**Figure 1:**
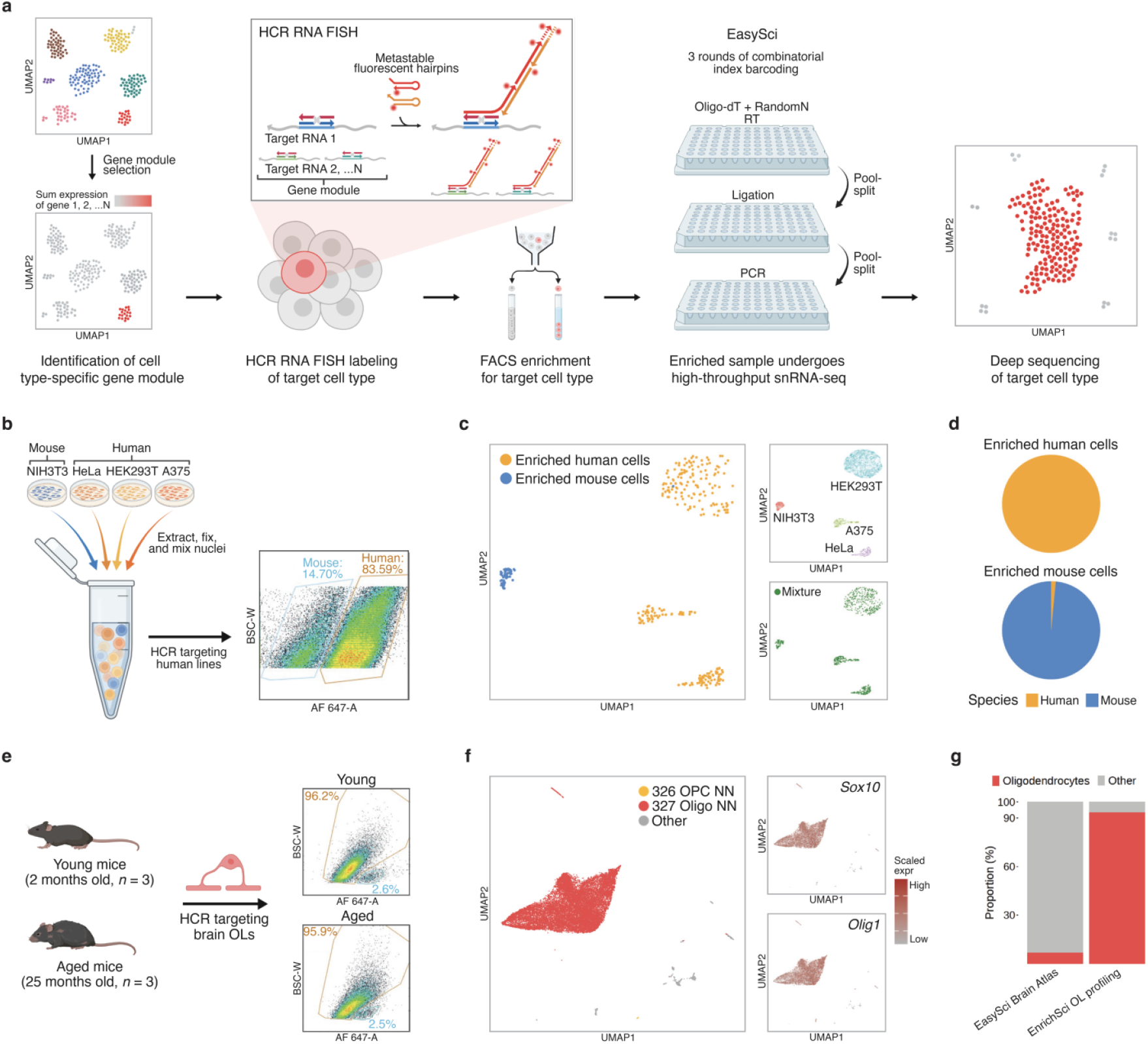
EnrichSci enables transcript-guided profiling of target cell types *in vitro* and *in vivo*. **a**, Scheme of EnrichSci workflow. **b**, Design of cell line mixture experiment (left) and FACS result (right). **c**, UMAP visualization of spike-in cell line controls (n = 8,338, top left), unenriched cell mixture (n = 909, bottom left), and enriched cells (n = 341, right). **d**, Pie charts showing proportion of cells from different species in enriched samples (Top: Enriched human cells, 100% assigned as human cells; Bottom: Enriched mouse cells, 98.5% assigned as mouse cells). **e**, Design of oligodendrocyte profiling experiment (left) and FACS result (right). **f**, UMAP visualization of mouse brain nuclei (n = 19,492) profiled by EnrichSci, colored by MapMyCells subclass name (left) and expression of oligodendrocyte lineage markers (right). **g**, Proportion of oligodendrocytes and non-oligodendrocytes in unenriched^1^ and enriched datasets.

As a proof of concept to demonstrate enriched cell profiling of EnrichSci, we extracted and fixed nuclei from mouse NIH3T3 cells and human HeLa, HEK293T, and A375 cells (**Fig. 1b**), mixed them to simulate a heterogeneous population, and performed HCR RNA FISH using probes against human-specific transcripts (*SCHLAP1, LIMCH1, PTMS, PDE4D, CAST, PDE3A, TPM4, IQGAP1, AKT3, RBMS3, SERPINE2, WWTR1, FTH1*) (**Supplementary Table 1)**. Next, we used FACS to separate fluorescent-positive (human) from fluorescent-negative (mouse) nuclei (**Fig. 1b**). Downstream EnrichSci profiling recapitulated the expected molecular states of each cell line (**Fig. 1c; Extended Data Fig. 2**), with nearly all cells (>98% of the fluorescent-positive and 100% of the fluorescent-negative cells) correctly mapping to their expected species compared with an unsorted spike-in control (**Fig. 1d**). We also confirmed that the HCR labeling and FACS steps did not compromise single-cell purity or RNA capture efficiency of the snRNA-seq workflow (**Extended Data Fig. 3**).

Building on our cell line validation, we next evaluated EnrichSci for analysis of specific cell populations from complex tissues by targeting oligodendrocytes in the mouse brain—a glial cell type critically involved in myelination and particularly susceptible to age-related degeneration^9^. Despite their importance, the full spectrum of oligodendrocyte subtypes and their gene expression dynamics across the entire gene body remains underexplored. To examine these details, we isolated nuclei from six male mice (three at 2 months and three at 25 months; **Fig. 1e**) and performed HCR labeling using probes against five oligodendrocyte-specific transcripts (*Mag, Galnt6, Opalin, Cyp2j12, Tnni1*; **Extended Data Fig. 4**; **Supplementary Table 1**). Notably, the proportion of fluorescent-positive nuclei was comparable between young and aged samples (**Fig. 1e**).

Following library preparation and sequencing, low-quality cells and doublets were removed (**Methods**), yielding 19,492 high-quality nuclei with a median of 3,566 unique transcripts (1,184 genes) per cell (**Extended Data Fig. 5a-b**). Cell identities were automatically annotated via the MapMyCells tool^10^ in conjunction with the Allen Brain Cell Atlas and confirmed by robust expression of oligodendrocyte markers *Mag* and *Plp1* (**Fig. 1f**). Compared with unbiased single-nuclei analysis of the whole brain that identified only 7% oligodendrocytes^1^, our targeted EnrichSci approach yielded an average of 93% oligodendrocytes—representing a 13-fold enrichment of the desired cell type (**Fig. 1g**).

To characterize the distinct subtypes of oligodendrocytes, we focused on cells annotated as oligodendrocyte precursor cells (OPCs) or oligodendrocytes and integrated data from different ages using the Seurat integration pipeline^11^. UMAP visualization revealed cellular states spanning the full oligodendrogenesis spectrum—OPCs (*Pdgfra+*), committed oligodendrocyte precursors (COPs, *Bcas1+*), newly formed oligodendrocytes (NFOL, *Prom1+*), myelin-forming oligodendrocytes (MFOL, *Slc9a3r2+*), and mature oligodendrocytes (MOLs, *Mog+*)^12^ (**Fig. 2a-b**; **Extended Data Fig. 5c**). Within the MOL population, we identified two distinct subtypes: MOL2, marked by *Klk6* and *Hopx*, which has been reported to be specific to hindbrain region^13,14^, and MOL5/6, characterized by expression of *Ptgds* and *Il33*, and found across broader brain regions^13,14^. Moreover, trajectory analysis showed that myelin-forming oligodendrocytes (MFOL) preferentially transition to MOL5/6 rather than to MOL2, suggesting that oligodendrocyte differentiation is impeded in the mouse hindbrain (**Fig. 2a-b**). This finding aligns with human single-cell studies reporting reduced oligodendrocyte differentiation in the cerebellum^15^.

**Figure 2.**
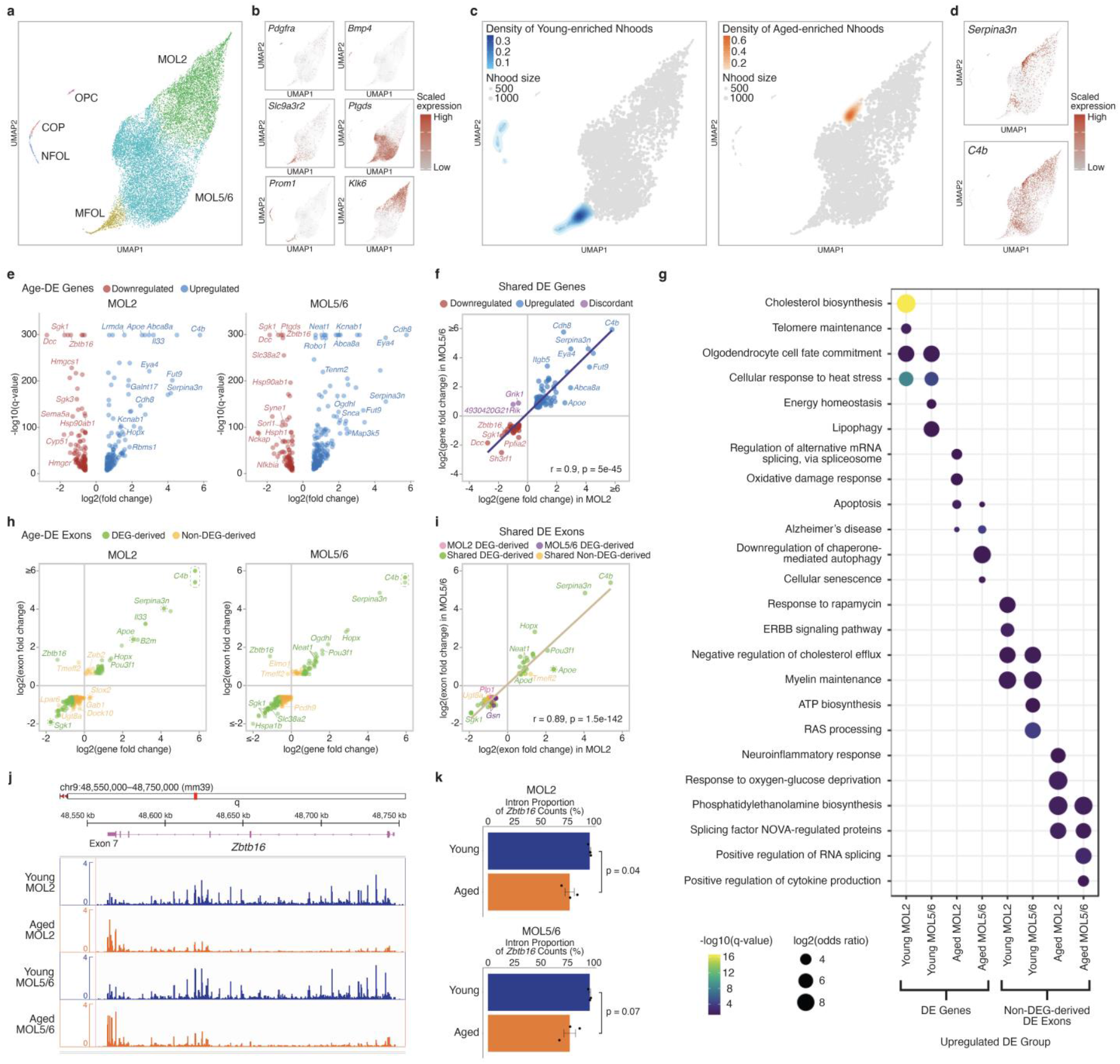
EnrichSci uncovers age-dependent transcriptome remodeling of brain oligodendrocyte subtypes at single-exon resolution. **a-b**, UMAP visualization of oligodendrocyte lineage nuclei (n = 18,154) profiled by EnrichSci, colored by subtype (a) or the expression of subtype-specific markers (b). **c**, UMAP visualization of cell neighborhoods, overlaid with density of Young-enriched (left) and Aged-enriched (right) neighborhoods. **d**, UMAP visualization of oligodendrocyte lineage nuclei, colored by expression of reactive oligodendrocyte markers (right). **e**, Volcano plots of differentially expressed (DE) genes between young and aged MOL2 (left) and MOL5/6 (right). Upregulated genes are shown in blue, downregulated genes in red. **f**, Scatter plot comparing log_2_ fold changes for the DE genes (DEGs) shared by both MOL subtypes. Points are colored by direction of change (upregulated, downregulated, or discordant between subtypes). **g**, Dot plot showing significantly enriched pathways for DEGs (left) and for DE exons (DEEs) not captured by DEGs (right). Results are stratified by subtype (MOL2 vs. MOL5/6) and age (young vs. aged). Dot size represents log_2_(odds ratio) and color indicates -log_10_(q-value). **h**, Scatter plots of DEEs in MOL2 (left) and MOL5/6 (right). Exons derived from DEGs are shown in green, non-DEG-derived exons in orange. **i**, Scatter plot comparing log_2_ fold changes for the DEEs shared by both MOL subtypes. Points are colored by DE status of the parent gene. **j**, Genome browser tracks showing aggregated read coverage across the *Zbtb16* gene body in young versus aged MOL2 and MOL5/6. **k**, Bar plots showing the proportion of *Zbtb16* reads mapping to intronic regions in young and aged MOL2 (top) and MOL5/6 (bottom). Individual mouse replicates are overlaid as points, with p-values calculated using an unpaired two-sided t-test.

To evaluate the effect of aging on cellular states, we used the Milo pipeline^16^ to cluster cells into transcriptionally similar neighborhoods and conducted differential abundance testing between young and aged samples. This revealed highly heterogeneous responses to aging across different cell states. Notably, intermediate precursor populations—including COPs, NFOL and MFOL—were significantly depleted in aged brains (**Fig. 2c**), reflecting impaired oligodendrocyte differentiation consistent with prior studies^1,15^. Although the overall mature oligodendrocyte pool remained relatively stable, we detected a distinct aging-associated MOL subpopulation marked by reactive genes such as *C4b* and *Serpina3n* (**Fig. 2d**). The concurrent loss of precursor cells and expansion of reactive mature oligodendrocytes aligns with observations from whole-brain single-cell analyses^1,14,17^, validating the capability of EnrichSci to dissect cellular subtypes and cell state transitions within targeted cell lineages.

We then examined how aging alters the transcriptomes of the two MOL subtypes. Using differential expression analysis (FDR < 0.05, fold change ≥ 1.5), we found 251 genes significantly dysregulated in MOL2 and 294 in MOL5/6 (**Fig. 2e, Supplementary Table 2**). Despite the distinct molecular profiles and regional distributions between the two subtypes, their aging-associated gene expression changes were remarkably concordant: the set of 119 DE genes identified in both subtypes showed highly correlated expression shifts (Pearson r = 0.9, p-value = 5e-45, **Fig. 2f**), indicating a global transcriptional remodeling of oligodendrocytes in aging. Common up-regulated genes included those involved in apoptosis (*e*.*g*., *Map3k5; Pik3r3, Mapk10* in MOL2; *Itpr1, Nfkb1* in MOL5/6^18^) and Alzheimer’s disease (*e*.*g*., *Apoe, Map3k5, Insr, Plcb4*^*18*^), whereas genes involved in heat stress response (*e*.*g*., *Hsph1, Hspa4l, Hsp90aa1, Hsp90ab1*^*18*^) and oligodendrocyte cell fate commitment (*e*.*g*., *Olig2*^*18*^) were down-regulated (**Fig. 2g**). We also observed reduced levels of mineralocorticoid targets such as *Sgk1* and *Sgk3*—key regulators of calcium channel activity and glucose uptake—and a broad down-regulation of myelination genes (*e*.*g*., *Mog, Plp1*) (**Extended Data Fig. 6**), matching prior reports of age-related myelin loss^17^. In addition to these shared signatures, each subtype exhibited unique aging pathways. Aged MOL2 specifically downregulates genes involved in cholesterol biosynthetic processes (*e*.*g*., *Hmgcs1, Hmgcr, Cyp51, Msmo1, Sqle, Sc5d, Dhcr7, Fdft1*^*18*^) and telomere maintenance (*e*.*g*., *Zfp827*^*18*^) and upregulates genes linked to oxidative damage response (*e*.*g*., *Mapk10*^*18*^) and spliceosome-mediated alternative mRNA splicing (*e*.*g*., *Celf2, Nova1*^*18*^). Meanwhile, aged MOL5/6 showed selective downregulation of genes involved in energy homeostasis (*e*.*g*., *Sorl1, Ubb*^*18*^) while upregulating genes associated with cellular senescence (*e*.*g*., *Itpr1, Atr, Foxo1, Nfkb1*^*18*^) (**Fig. 2g**).

EnrichSci uniquely enables full gene-body coverage in targeted cell populations, thereby allowing exon-level expression analysis. By applying the same differential expression framework at the exon scale (FDR of 0.05, fold change ≥ 1.5), we identified 261 and 254 differentially expressed exons (DEEs) in MOL2 and MOL5/6, respectively (**Fig. 2h, Supplementary Table 3**). Of these, 103 DEEs were shared between both subtypes and exhibited highly coordinated aging dynamics (Pearson r = 0.89, p = 1.5e-142, **Fig. 2i**). Although most DEEs exhibited similar dynamics as their parent genes—for example, exons from *C4b* and *Serpina3n* were up-regulated, while exons from *Sgk1* and *Ugt8a* were down-regulated—a substantial fraction (42.5% of DEEs in MOL2; 40.1% in MOL5/6; 30.1% of shared DEEs) did not overlap with DEGs (**Extended Data Fig. 7a-b**). These non-DEG-derived DEEs uncover pathway perturbations invisible to traditional gene-level analyses (**Fig. 2g**). For instance, aged MOL5/6 exhibited enrichment of exons in genes associated with positive regulation of RNA splicing (*e*.*g*., *Pik3r1, Srsf5*^*18*^) and cytokine production (*e*.*g*., *Sptbn1, Pik3r1*^*18*^). In contrast, aged MOL2 displayed increased exon usage in genes linked to neuroinflammatory response (*e*.*g*., *Zeb2*^*18*^) and a down-regulation of the ERBB-signaling exons, a pathway previously implicated in the regulation of oligodendrocyte differentiation (**Fig. 2c**).

Finally, some DEEs showed inverse trends from the genes they derive from. One striking example is *Zbtb16*, a transcription factor essential for oligodendrocyte maturation whose knockout impairs social cognition and myelination in the prefrontal cortex^19^. Although overall *Zbtb16* gene expression significantly declined with age, exon 7 near its 3′ end was substantially more included in aged MOL2 and MOL5/6 (**Fig. 2j**). This discordance reflects a shift in splicing dynamics—young MOL2 and MOL5/6 showed high intronic read fractions for *Zbtb16* (96.1% and 96.1%, respectively), which dropped to 77.1% and 77.3% in aged cells (**Fig. 2k**). Together, these exon-level changes underscore post-transcriptional regulation as an additional layer of cell-state dynamics during aging.

In summary, while large-scale single-cell atlas projects have cataloged numerous aging- and disease-associated cell states^1,2,20,21^, many of these populations remain poorly characterized in terms of subtype diversity and detailed molecular signatures. To overcome this limitation, we introduce EnrichSci—an antibody-free, HCR-based single-nucleus sequencing method for targeted profiling of specific cell types within complex tissues. Unlike previous approaches^3–5^, EnrichSci represents the first single-cell combinatorial indexing strategy that combines efficient enrichment of target cell types and highly scalable single-cell analysis (from thousands to millions of cells^1^) with genome-wide RNA coverage across full gene bodies, and it is easily implemented with standard laboratory equipment. This platform enables the simultaneous detection of both gene-level and exon-level aging signatures in rare cell types such as oligodendrocyte subtypes, revealing post-transcriptional regulatory changes that are missed by conventional analyses. Looking ahead, EnrichSci can be readily expanded to incorporate additional molecular layers (*e*.*g*., chromatin accessibility^22^, DNA methylation^23^) and genetic perturbations (*e*.*g*., CRISPR^24^), providing a versatile platform to dissect molecular dynamics and identify genetic drivers in aging- and disease-associated cell types *in vivo*.

## Endnotes

## Acknowledgments

We thank all members of the Cao Lab for helpful discussions and feedback. We thank the Tissue Culture facility of the University of California, Berkeley, for the NIH/3T3 cell line, and the Scott Keeney Lab at Memorial Sloan Kettering Cancer Center for the HEK293T cell line. We also thank members of the Information Technology and High-Performance Computing team at Rockefeller University, especially J. Banfelder and B. Jayaraman, for their great support. We acknowledge that the research resulting in this publication was supported, in part, by The G. Harold and Leila Y. Mathers Charitable Foundation.

## Author contributions

J.C. and W.Z. conceptualized and supervised the project. A.L. performed all experiments, including technique development and optimization, with input from other co-authors. A.L. performed all computational analyses, with input from other co-authors. J.C., W.Z., and A.L. wrote the manuscript with input and biological insight from all co-authors.

## Data and materials availability

Raw FASTQ files, processed count matrices, cell metadata, and gene metadata can be downloaded from NCBI GEO under accession number GSE295135 (GEO reviewer token: **gbqbiwggpbelhsz**).

## Supplementary Figures

**Extended Data Fig. 1.**
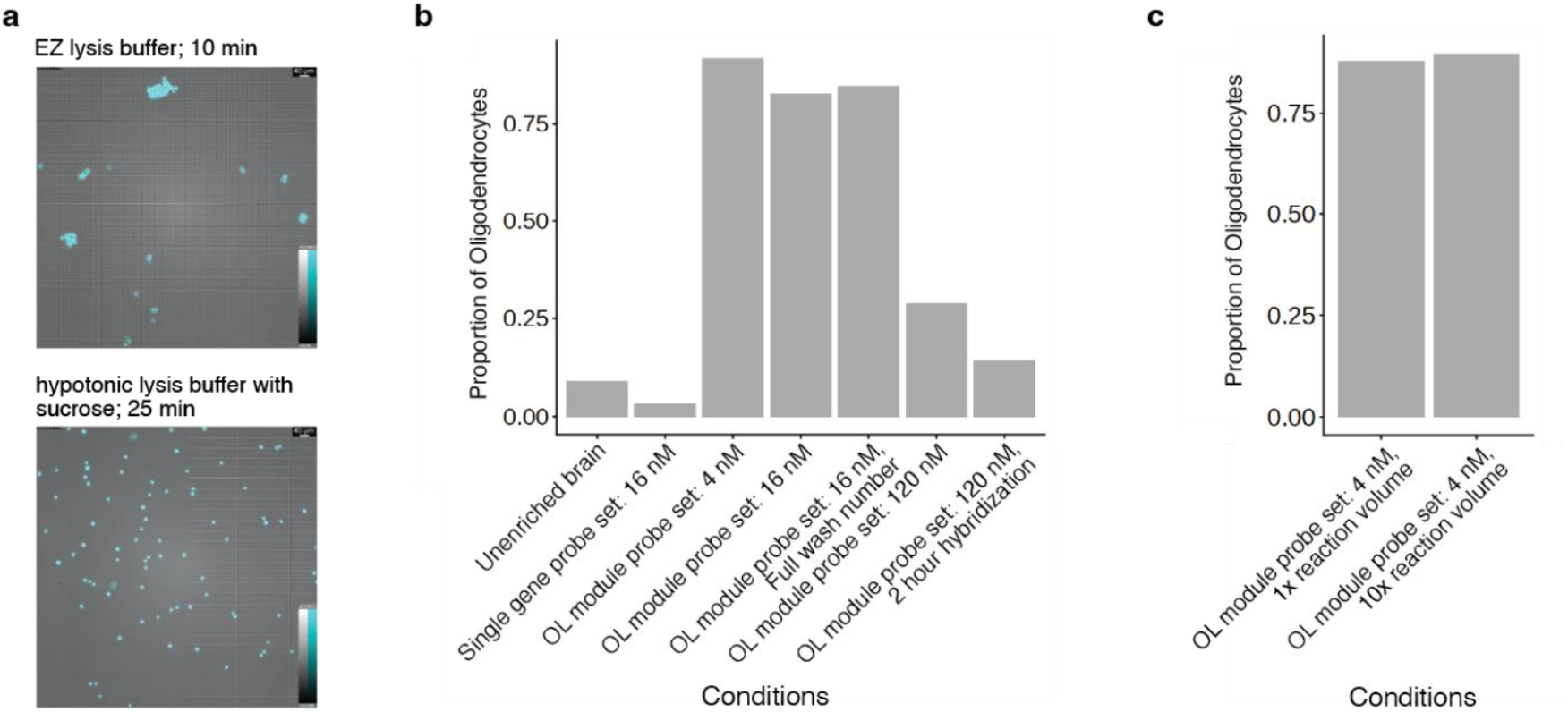
Optimization of EnrichSci performance. **a**, Comparison of nuclei extraction buffers for A375 cells. Nuclei prepared in a hypotonic sucrose buffer (bottom) exhibit markedly reduced clumping and higher recovery than those processed with the commercial EZ Lysis Buffer (top). **b**, Evaluation of HCR RNA FISH conditions for oligodendrocyte enrichment from mouse brain. We varied probe design (single -gene *Sox10* vs. multi-gene oligodendrocyte modules), probe concentration (4, 16 or 120 nM), wash stringency (standard vs. reduced number of washes) and hybridization time (2 h vs. overnight), and assessed enrichment efficiency by single-cell sequencing. The optimal settings—multi-gene probe modules at 4 nM, reduced wash steps, and overnight hybridization—yielded the highest recovery of target cells. **c**, Comparison of HCR reaction volumes (1× vs. 10×) for large-scale processing of mouse brain nuclei. Both volumes produced comparable enrichment efficiencies.

**Extended Data Fig. 2.**
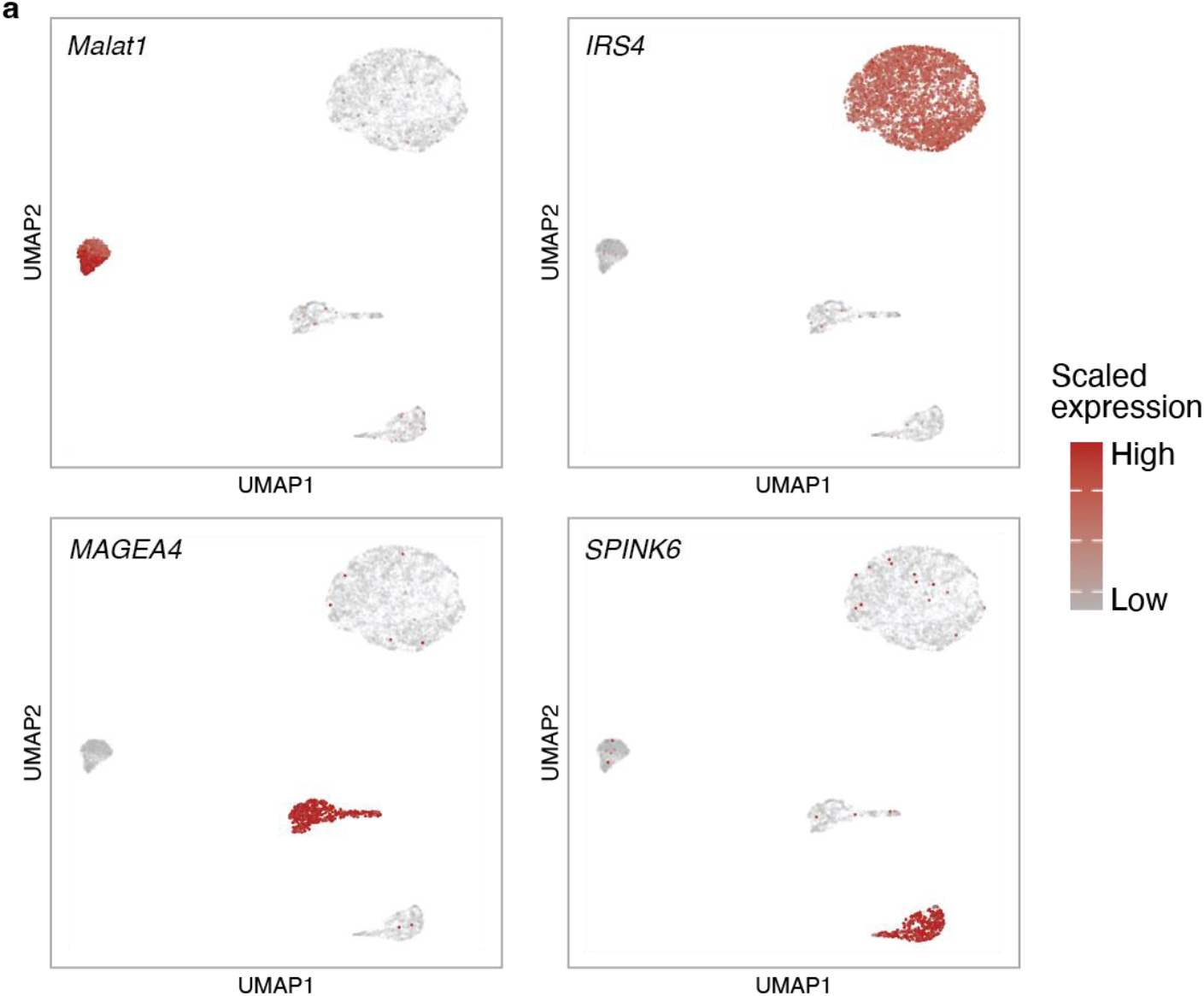
EnrichSci recapitulates expected molecular states of cell lines profiled. UMAP visualization of single-cell transcriptomes that include individual cell line spike-ins, unenriched mixture, and enriched samples profiled by EnrichSci, colored by the normalized and scaled expression of cell line-specific gene markers.

**Extended Data Fig. 3.**
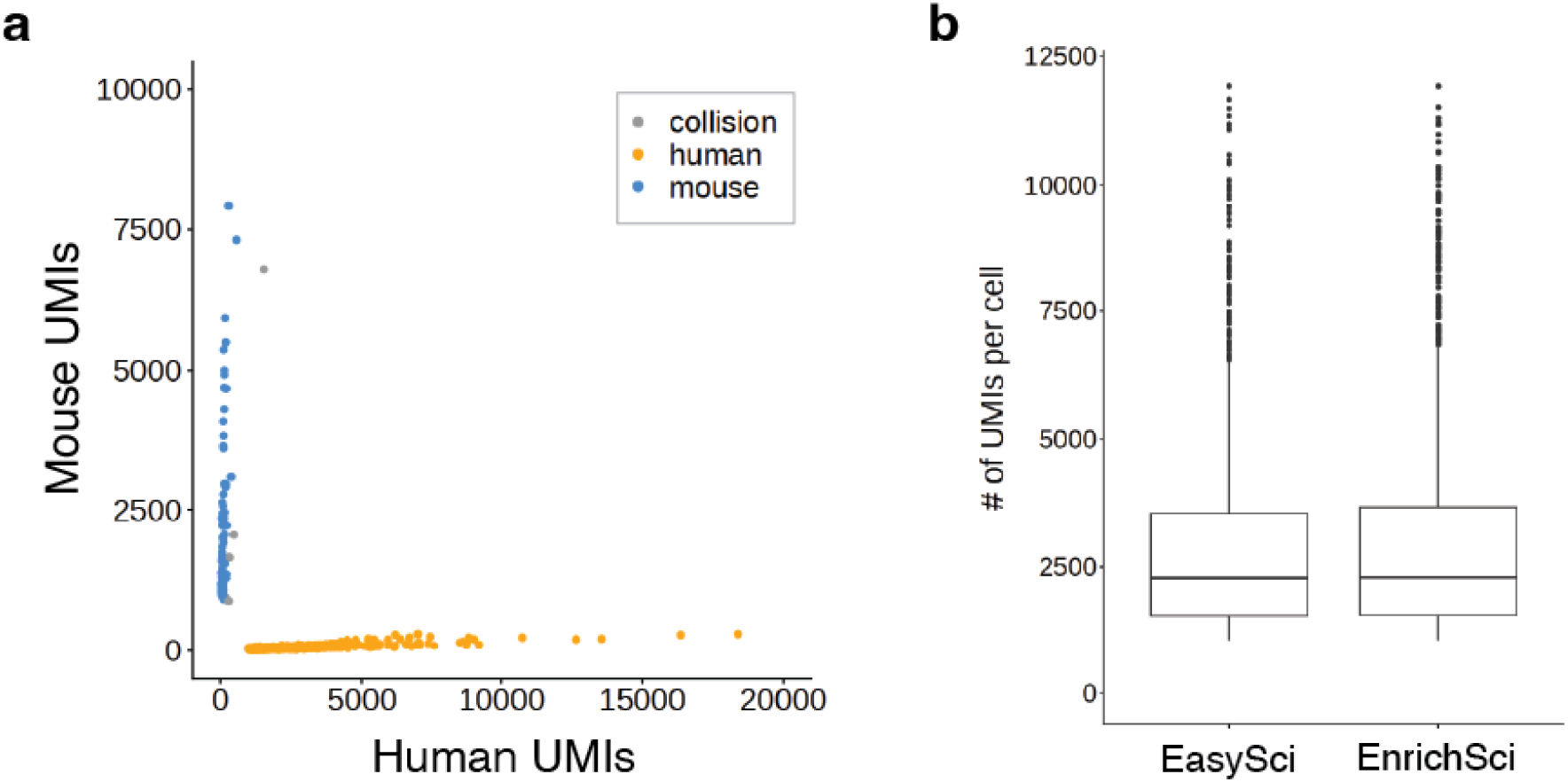
HCR labeling does not compromise single-cell purity or RNA capture efficiency. **a**, Scatter plot of mouse and human unique molecular identifier (UMI) counts from the cell line mixture profiled by EnrichSci. Blue, inferred mouse cells (n = 85). Orange, inferred human cells (n = 814). Gray, collisions (n = 10). **b**, Boxplot showing UMI count distribution of cells profiled by EasySci (n = 1,535; median = 2,296) and EnrichSci (n = 1,571; median = 2,331).

**Extended Data Fig. 4.**
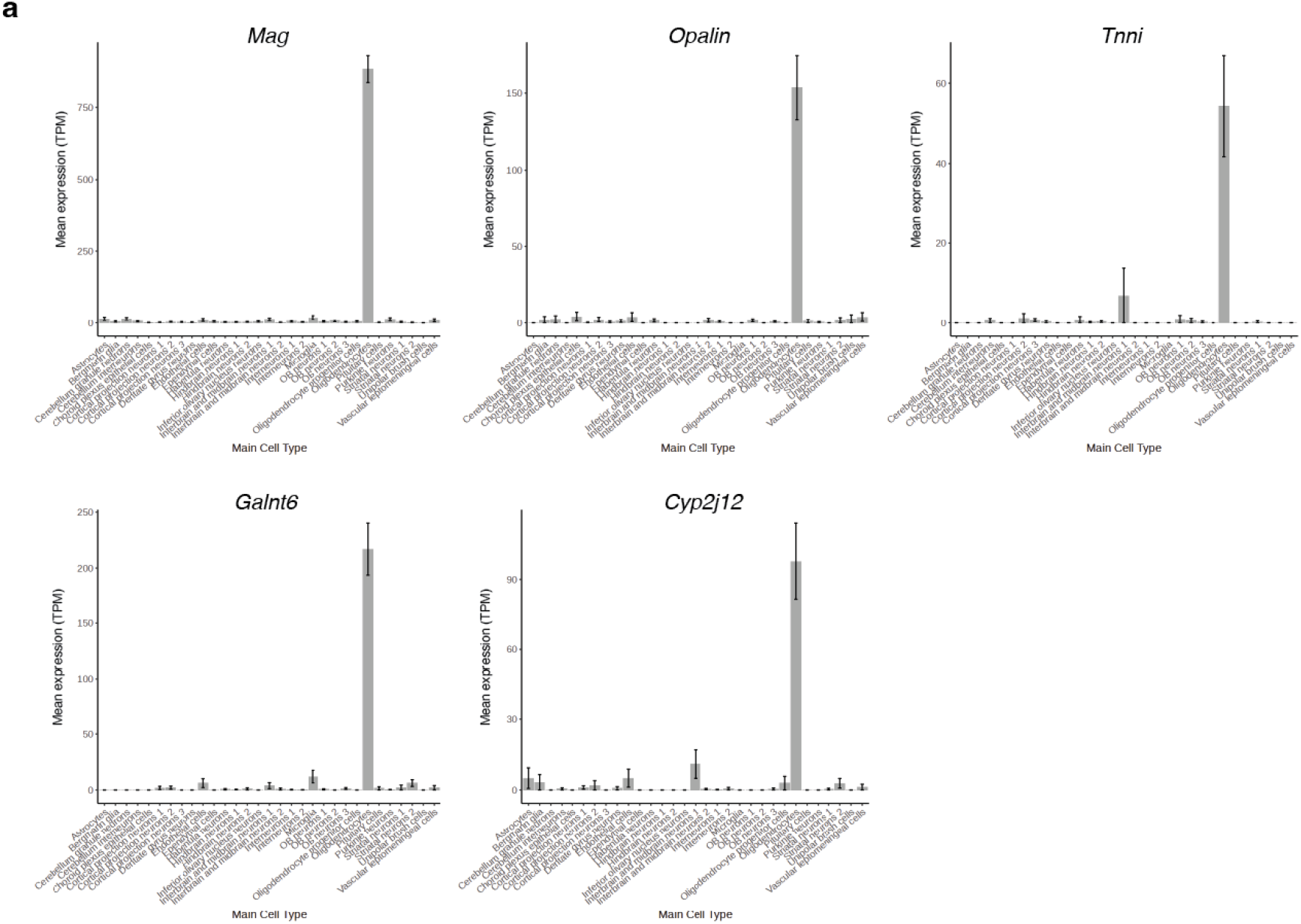
Selection of oligodendrocyte-specific genes. Barplots showing expression of oligodendrocyte-specific genes across main cell types in the mouse brain^1^. Expression is normalized to transcripts per million (TPM) and averaged across replicates.

**Extended Data Fig. 5:**
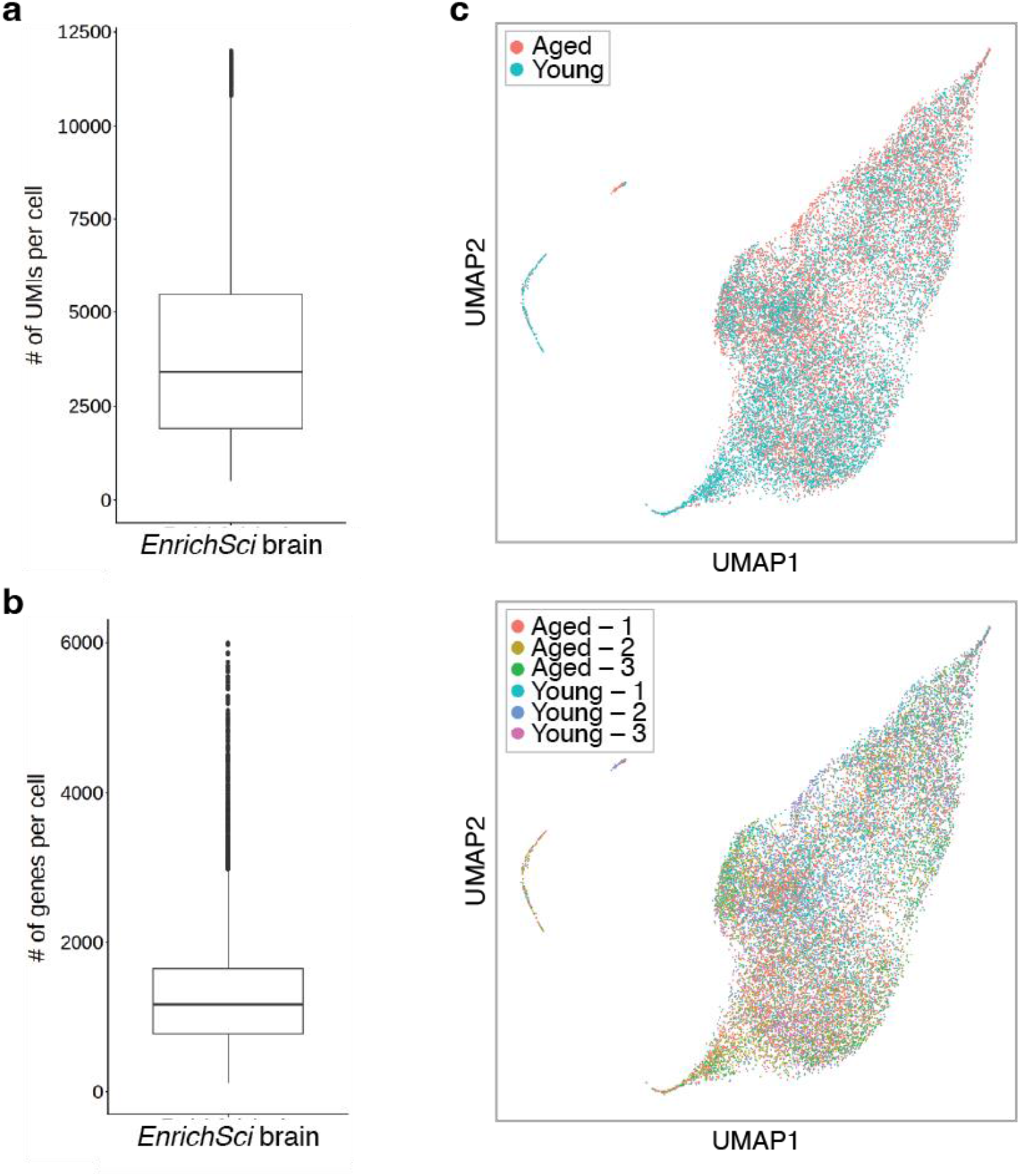
EnrichSci recovered single-cell transcriptome profiles of mouse brain nuclei with high signals and sample consistency. **a**, Boxplot showing UMI count distribution of oligodendrocyte-enriched mouse brain nuclei profiled by EnrichSci. **b**, Boxplot showing gene count distribution of oligodendrocyte-enriched mouse brain nuclei profiled by EnrichSci. **c**, UMAP visualization of oligodendrocyte lineage nuclei (n = 18,154) profiled by EnrichSci, colored by age group (top) and individual replicate (bottom).

**Extended Data Fig. 6:**
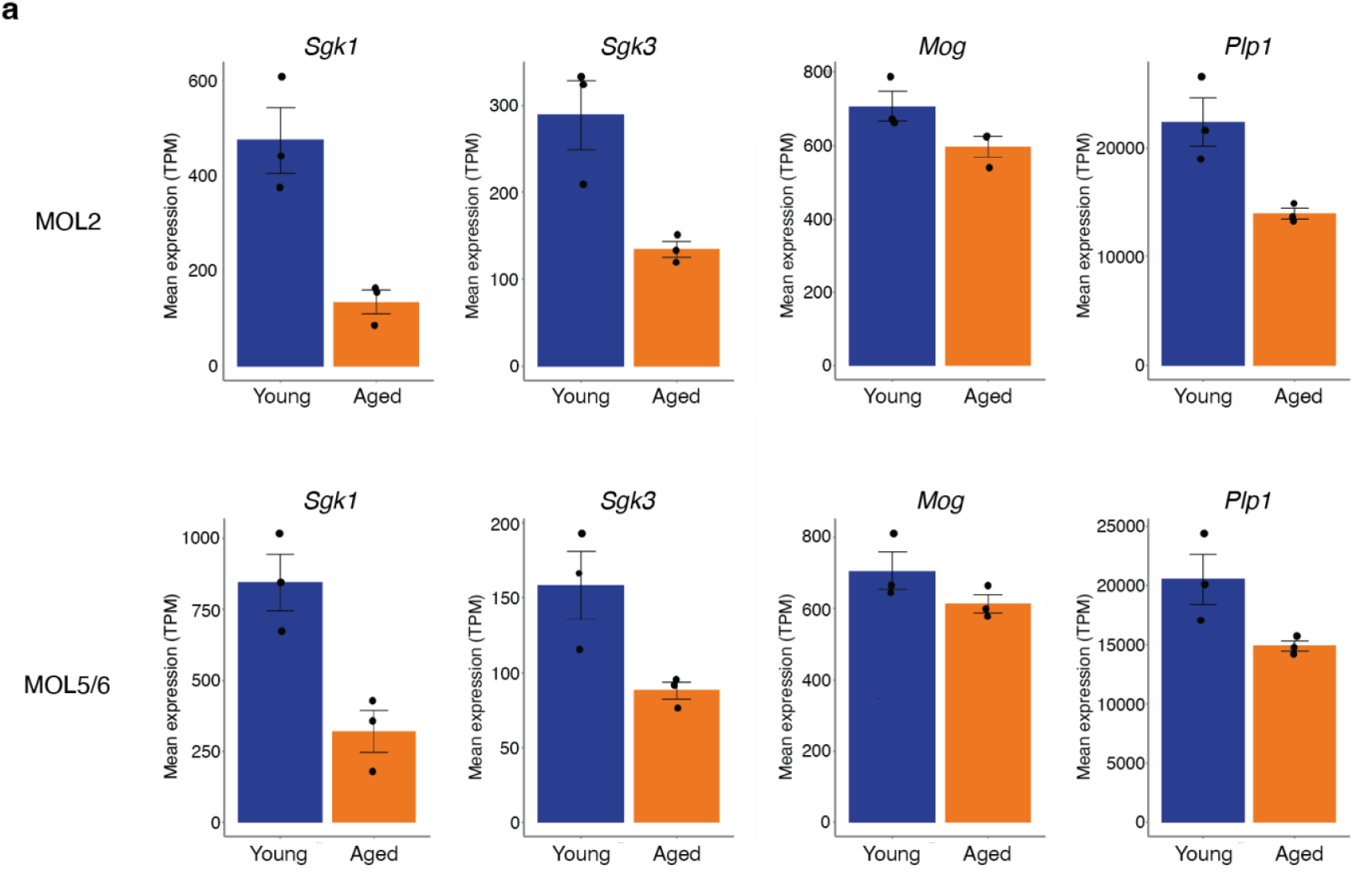
Mature oligodendrocyte subtypes exhibit decreased expression of key oligodendrocyte genes with age. **a**, Barplots showing expression of *Sgk1, Sgk3, Mog*, and *Plp1* in young and aged MOL2 (top) and MOL5/6 (bottom). Expression is normalized to transcripts per million (TPM) and averaged across three replicates.

**Extended Data Fig. 7:**
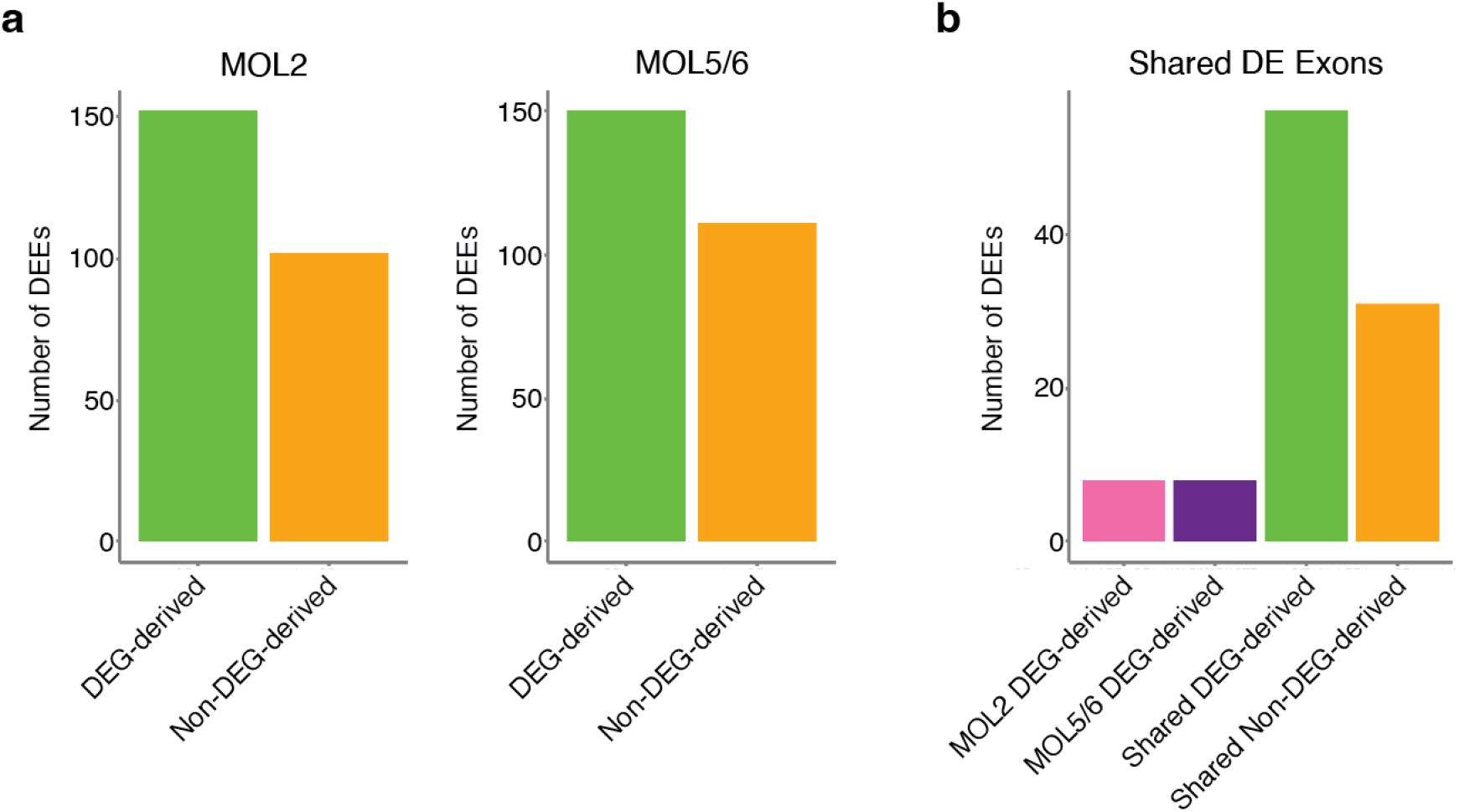
EnrichSci identifies substantial numbers of differentially expressed (DE) exons derived from genes that are not DE. **a**, Barplots showing the number of DE exons (DEEs) that derive from DE genes (DEGs) and non-DEGs in MOL2 (left) and MOL5/6 (right). **b**, Barplot showing number of shared DEEs that derive from subtype-specific DEGs, shared DEGs, and shared non-DEGs.

## Materials and Methods

### Cell Culture

HEK293T (gift from Scott Keeney Lab, Memorial Sloan Kettering Cancer Center), A375, HeLa, and NIH/3T3 (gift from Tissue Culture facility of the University of California, Berkeley) cell lines were cultured in 10 cm dishes at 37°C with 5% CO_2_ in high glucose DMEM (Gibco, 11965-118) supplemented with 10% Fetal Bovine Serum (Sigma-Aldrich, F4135) and 1% penicillin-streptomycin (Gibco, 15140-122).

### Animals

C57BL/6 wild-type mice at two months (n = 3) and twenty-five months (n = 3) were obtained from The Jackson Laboratory. Mice were housed socially and maintained on a regular 12h/12h day/night cycle. Euthanization and tissue collection were performed around the same period of time during the day to control for circadian effects. Mice were euthanized utilizing inhalation of carbon dioxide (CO2), followed by cervical dislocation, prior to tissue dissection. All animal procedures were in accordance with institutional, state, and government regulations and approved under the IACUC protocol 24012-H.

### Tissue collection and nuclei isolation

After euthanization, whole brains were extracted from mice, immediately snap-frozen in liquid nitrogen, and stored at -80°C for further usage. For nuclei isolations, thawed brains were diced into fine pieces using carbon steel razor blades (VWR, 100491-872) in a 6 cm dish containing 1 mL hypotonic lysis buffer with sucrose (7.68 mM Na2HPO4·2H2O, 4.49 mM NaH_2_PO_4_·H_2_O, 1.76 mM KH_2_PO_4_, 2.68 mM KCl, 10.27 mM NaCl, 3 mM MgCl2, 0.33 M sucrose, 0.025% IEGPAL, 1% DEPC), transferred to a 15 mL tube containing 14 mL of hypotonic lysis buffer with sucrose for a 25 minute incubation while rotating at 4°C, and then homogenized through 40 µm cell strainers (Ward’s Science, 470236-276). Extracted nuclei were then pelleted, fixed in 10 mL 0.4% formaldehyde while rotating for 15 minutes at 4°C, and washed twice with Nuclei Suspension Buffer (NSB) (10 mM Tris-HCl pH 7.5 (VWR, 97062-936), 10 mM NaCl (VWR, 97062-858), 3 mM MgCl2 (VWR, 97062-848) supplemented with 0.1% SUPERase-In RNase Inhibitor (Thermo Fisher Scientific, AM2696), 1% BSA (NEB, B9200S), and 0.1% Tween-20 (Sigma, P9416-100ML). Nuclei were counted and resuspended at 1 million nuclei/mL in NSB with 10% DMSO, cryopreserved at -80°C using a controlled-rate freezing container (Corning, 07-210-009), and stored at -80°C until usage.

### EnrichSci HCR and FACS enrichment protocol

HCR was performed largely according to the Molecular Instruments (MI) “*HCR RNA flow cytometry protocol for mammalian cells in suspension*” protocol (version dated 2023-02-13), with specific optimizations made for compatibility with nuclei, improved nuclei recovery, and optimal signal-to-noise. Of note, the number of wash steps during the detection and amplification stages was significantly reduced by 7 washes from the original MI protocol. All centrifugation steps to pellet nuclei were performed at 1,200g (4°C) for 5 minutes unless otherwise specified. All incubations were performed using a rotating mixer. Standard 1x reaction volumes are described below. For the oligodendrocyte-enriched mouse brain samples, all steps prior to pre-amplification were scaled up to 5× reaction volumes and carried out in 15 mL tubes. At the pre-amplification step, the reaction was condensed back to the standard 1x volume.

As described above, extracted nuclei were fixed with 0.4% formaldehyde, in contrast to the 4% formaldehyde fixation and ethanol permeabilization used for whole cells in the MI protocol, and stored at -80°C until usage. Frozen, fixed nuclei from each sample were thawed from -80°C in a 37°C water bath, pelleted, and resuspended in 400 μL of MI probe hybridization buffer. After a 30-minute pre-hybridization incubation at 37°C, 100 μL of probe solution (prepared by adding 1 μL of probe stock at 2 μM per probe to 99 μL of hybridization buffer) was added to each sample to achieve a final concentration of 4 nM, and samples were incubated overnight at 37°C.

The following day, 500 uL of MI probe wash buffer was added to each sample before pelleting at 1,600g (4°C) for 5 minutes. Supernatant was removed, and pellets were resuspended in 1 mL of probe wash buffer and incubated at 37°C for 10 minutes. Each sample was then pelleted at 1,400g (4°C) for 5 minutes, resuspended in 1 mL of 5x SSCT (5x saline sodium citrate, 0.1% Tween-20), and incubated at room temperature for 5 minutes. Samples were pelleted, resuspended in 150 μL of MI amplification buffer, and rotated at room temperature for 30 minutes in a pre-amplification incubation. After pre-amplification, 110 μL of amplifier solution (prepared by adding 5 μL each of MI amplifier h1 and h2 to 100 μL of amplification buffer) was added to each sample. Samples were incubated at room temperature overnight.

The following day, 1 mL of 5x SSCT was added to each sample before pelleting. Samples were washed again in 1 mL of 5x SSCT, then resuspended in NSB containing DAPI (Thermo Fisher Scientific, D1306) at a 1:50 dilution from a 0.25 mg/mL stock before proceeding to nuclei sorting. FACS was performed using a SH800 Cell Sorter with a 100 μm sorting chip (Sony, #LE-C3210). Nuclei were first gated to select DAPI-positive singlets, followed by gating for populations of interest based on Alexa Fluor 647 fluorescence from HCR. Sorting was carried out into 1.5 mL tubes pre-coated with NSB.

### EnrichSci snRNA-seq protocol

Sorted nuclei underwent combinatorial indexing-based sequencing library generation according to the EasySci^1^ protocol. In brief, sorted nuclei were first distributed into 96-well plates (Genesee Scientific, #24-302), each with well-specific oligo-dT and random hexamer primers that provided the first round of indexing via reverse transcription. After reverse transcription, nuclei were pooled, washed, and redistributed into new 96 -well plates for a second round of indexing via ligation. Finally, nuclei were pooled, washed, and redistributed into 8 -well PCR strips (only up to 8 strips were needed for the number of brain nuclei profiled in this study). Second-strand synthesis was performed and stored at -20°C overnight before purification the next day. After purification, tagmentation with Tn5 transposase was performed, and then the final round of indexing was achieved during the final indexed PCR. The PCR products were pooled and purified twice using AMPure XP Beads (Beckman Coulter, #A63882), first using a 0.8X volume and again using a 0.9X volume. Libraries were visualized by gel electrophoresis, and concentrations were determined using a Qubit fluorometer (Invitrogen, Q33231). All libraries were sequenced on the NextSeq 1000 platform (Illumina) using a 100-cycle kit (Read 1: 58 cycles, Read 2: 60 cycles, Index 1: 10 cycles, Index 2: 10 cycles). The cell line library was sequenced to ∼13,000 reads per cell, and the mouse brain library was sequenced to ∼37,000 reads/cell.

### EnrichSci data processing

Raw sequencing data were processed using the previously developed EasySci^1^ pipeline for read alignment and generation of gene and exon count matrices for snRNA-seq libraries. In brief, base calls were converted to FASTQ format and demultiplexed using Illumina bcl2fastq (v2.19.0.316), allowing up to one mismatch in barcode sequences (edit distance < 2). RT barcodes were corrected to their nearest valid barcode (edit distance <2), and reads with barcodes that could not be corrected (edit distance ≥ 2) were excluded. Adaptors and barcodes were trimmed, and Trim Galore^25^ (v0.4.1) was used to additionally remove poly(A) sequences and low-quality base calls. Using STAR^26^ (v2.5.2b), trimmed reads were aligned to a chimeric human and mouse genome (hg27/mm10) for the cell line mixture experiment and the mouse genome (mm39) for mouse brain profiling. After removal of PCR duplicates, which share a unique molecular identifier (UMI) sequence, RT barcode, and tagmentation site, reads are split into SAM files per cell. A custom EasySci^1^ script was used to quantify gene and exon expression per cell. To assign reads to genes, a read was counted if its aligned coordinates overlapped with annotated gene regions. If a read derived from the oligo-dT RT primer was ambiguous between multiple genes, it was assigned to the gene with the closest 3’ end. If a read did not initially map to a gene, the script also searched for potential gene assignments up to 1,000 bp upstream or on the opposite strand. After these steps, reads without assigned genes were discarded. A similar approach was performed to generate the exon count matrix.

### Cell filtering, clustering, and annotation for EnrichSci

Gene and exon expression matrices were constructed from the raw sequencing data as described above. In the cell line mixture experiment, cells with less than 1000 UMIs and 100 unique genes were discarded. For mouse brain profiling, cells with less than 500 UMIs and 100 unique genes were discarded.

To identify doublets, Scrublet^27^ (version 0.2.3) with Scanpy^28^ (v1.6.0) was applied to each gene count matrix using the following parameters: min_count = 3, min_cells = 3, vscore_percentile = 85, n_pc = 30, expected_doublet_rate = 0.06, sim_doublet_ratio = 2, n_neighbors = 30. Cells with doublet scores over 0.2 were annotated as doublets and discarded, along with any cells from doublet-derived sub-clusters. Finally, cells that passed initial filtering but appeared to be doublets, based on clustering results and expression of markers from multiple cell types, were also removed.

Using Seurat^11^ (v4.0.2), dimension reduction was performed on the data first by PCA using 30 components and then with UMAP before Louvain clustering. For the cell line mixture experiment, cluster identity was annotated using individual cell line spike-ins as a reference. For mouse brain profiling, we performed two independent experimental batches. Although both batches were integrated to define cell-state identities, downstream gene-expression analyses were restricted to the second batch—comprising three biological replicates per age group versus only two aged replicates in the first batch. Initial cell type annotations were generated using the Allen Institute for Brain Science MapMyCells tool^10^. We then removed all cells that did not have the class_name ‘31 OPC-Oligo’ to filter for only oligodendrocyte lineage cells (n = 18,250). For higher resolution clustering of these cells, we used Seurat^11^ to integrate oligodendrocyte lineage cells from the two independent experiment batches, by regressing out the effect of experimental batch and age group. Following integration, we performed dimension reduction and clustering, revealing shared and divergent cell states between the two age groups. For downstream analyses, only the second batch with all three replicate samples per age group was used. Oligodendrocyte lineage subtypes were annotated based on the previous MapMyCells^10^ annotations along with expression of cell type-specific markers (**Fig. 2e**).

### Cell population dynamics analysis

To assess the effects of aging on cell population dynamics within the oligodendrocyte lineage, we applied miloR^16^ (v1.3.1), a single-cell differential abundance testing framework using k-nearest neighbor (KNN) graphs. We first constructed a KNN graph on the UMAP space using the buildGraph() function with k = 60. Cell neighborhoods were defined with makeNhoods(), and cell counts per sample within each neighborhood were computed using countCells(). Differential abundance testing was performed using testNhoods(), with significance assessed at a spatial FDR threshold of 0.05. To visualize differential abundance neighborhoods, we initially tried the plotNhoodGraphDA() function, but it is sensitive to local noise and does not easily show the density of specific conditions. To better visualize age-related cell population shifts, we first expanded each neighborhood by including all cells within a fixed radius of its UMAP coordinates. We then calculated the proportion of young and aged cells in each neighborhood, labeling them as *Young-enriched* or *Aged-enriched* if one age group comprised more than 70% of the cells. These expanded neighborhoods were more robust to local variability and better captured broader population shifts. Finally, to visualize the distribution of age group-enriched neighborhoods, we used ggplot2 to overlay density contours on a UMAP plot of the cell neighborhoods.

### Differential expression analysis

To identify differentially expressed (DE) genes and exons between young and aged mice in each of the two mature oligodendrocyte subtypes, we employed the likelihood ratio test to identify genes and exons significantly associated with a specific population, using the differentialGeneTest() function in Monocle2 (version 2.28.0^29^). Filtering first for DE genes, we used the following cutoffs: FDR < 0.05, fold change > 1.5, and transcripts per million (TPM) in the max condition > 25. To filter for DE exons, we used the same FDR and fold change cutoffs but lowered the TPM threshold (in the max condition) to > 10, accounting for the lower expression levels of individual exons. To classify DE exons as either DEG-derived or non-DEG-derived, we re-filtered genes using the DE exon TPM cutoff (> 10) instead of > 25. Using the same cutoffs as DE exon filtering thus allowed us to determine that under the same criteria, the parent genes of non-DEG-derived DE exons were only detectable at the exonic and not the gene level.

### Gene set enrichment analysis

To identify enriched pathways associated with DE genes and exons, we used g:Profiler^18^ (version e112_eg59_p19_25aa4782) with Benjamini-Hochberg FDR < 0.05 applied. DE genes and non-DEG-derived DE exons for each mature oligodendrocyte subtype and age group were separately analyzed. DE gene inputs consisted of unranked mouse gene symbols, and DE exon inputs used the corresponding parent gene symbols. We queried the following pathway databases: GO Biological Process (2024-10-7 release), KEGG pathways (2024-01-22 release), Reactome pathways (2025-02-03 release), and WikiPathways (2025-01-10 release). For visualization, we manually selected biologically relevant pathways for DE genes and exons in each age group and mature oligodendrocyte subtype and plotted them using custom ggplot2-based scripts.

## Supplementary Tables (provided as Microsoft Excel files)

**Supplementary Table 1:** Probe and primer sequences used in EnrichSci experiments. Probe sequences are included for both the cell line mixture and the oligodendrocyte-enriched brain profiling experiments. Full primer sequences, well positions, and barcode sequences of the oligo-dT RT primers, random hexamer RT primers, ligation primers, and P5/P7 primers are also included.

**Supplementary Table 2:** Aging differentially expressed genes across mature oligodendrocyte (MOL) subtypes. The MOL subtype, the highest expressing age group, gene name, gene ID, q-value, log_2_ fold change in aged versus young mice, and TPM are included.

**Supplementary Table 3:** Aging differentially expressed exons across mature oligodendrocyte subtypes. The MOL subtype, the highest expressing age group, the gene name, the exon ID, q-value, log_2_ fold change in aged versus young mice of the exon, log_2_ fold change of the parent gene, TPM, and DE status of the parent gene are included.

